# Obesogenic diet induces circuit-specific memory deficits in mice

**DOI:** 10.1101/2022.06.20.496841

**Authors:** Ioannis Bakoyiannis, Eva-Gunnel Ducourneau, Matéo N’diaye, Alice Fermigier, Céline Ducroix-Crepy, Clémentine Bosch-Bouju, Etienne Coutureau, Pierre Trifilieff, Guillaume Ferreira

**Author notes:** co-first author (Ioannis Bakoyiannis, Eva-Gunnel Ducourneau).

## Abstract

Obesity is associated with neurocognitive dysfunction, including memory deficits. This is particularly worrisome during adolescence, which represents a crucial period for maturation of brain structures, such as the hippocampus which are crucial for cognition. In rodent models, we recently reported that memory impairments induced by obesogenic high-fat diet (HFD) intake during the periadolescent period can be reversed by chemogenetic manipulation of the ventral hippocampus (vHPC). Here, we used an intersectional viral approach in HFD-fed male mice to chemogenetically inactivate specific vHPC efferent pathways to nucleus accumbens or medial prefrontal cortex during memory tasks. We first demonstrated that HFD enhanced activation of both pathways after training and that our chemogenetic approach was effective in normalising this activation. Inactivation of the vHPC-nucleus accumbens pathway rescued HFD-induced deficits in recognition but not location memory. Conversely, inactivation of the vHPC-medial prefrontal cortex pathway restored location but not recognition memory impairments produced by HFD. Either pathway manipulation did not affect exploration, locomotion or anxiety-like behaviour. These findings suggest that HFD intake throughout adolescence impairs different types of memory through overactivation of specific hippocampal efferent pathways and that targeting these overactive pathways has therapeutic potential.

## Introduction

Obesity is a chronic disease critically affecting public health that has been rising tremendously over the last years. Primarily due to an overconsumption of energy-dense food combined with a sedentary lifestyle, obesity is associated with several comorbidities including cardiovascular and metabolic diseases (Carbone et al., 2013), but also cognitive disorders (Francis and Stevenson, 2013; Sui and Pasco, 2020; Wang et al., 2014). In particular, memory deficits have been previously reported in obese adults (Francis and Stevenson, 2013; Martin and Davidson, 2014; Sellbom and Gunstad, 2012; Yeomans, 2017), but also in obese adolescents (Khan et al., 2015; Nyaradi et al., 2014; Øverby et al., 2013). This is worrisome because the prevalence of obesity has risen during adolescence (Ogden et al., 2016) which is a crucial period for brain development (Andersen, 2003; Spear, 2000). It is therefore timely to identify the mechanisms by which adolescent obesity impairs cognitive functions.

Using animal models, we and others have extensively demonstrated the higher vulnerability of adolescence to the effects of obesogenic diet on hippocampal (HPC) function and HPC-dependent memory as compared to adulthood (Boitard et al., 2014, 2012; Del Olmo and Ruiz-Gayo, 2018; Glushchak et al., 2021; Khazen et al., 2019; Morin et al., 2017; Murray and Chen, 2019; Tsan et al., 2021; Valladolid-Acebes et al., 2013). However, increasing evidence suggest that, rather than homogeneously altering neuronal function, discrete neuronal subpopulations and pathways are particularly vulnerable to dietary manipulations(Berland et al., 2020; Ducrocq et al., 2020). Accordingly, few studies reported some effects of adolescent obesity on specific functional circuits in humans (Vega-Torres et al., 2018) and animal models (Moreno-Castilla et al., 2018). These findings made us hypothesize that deficits in distinct HPC-dependent memory could result from alterations of discrete HPC pathways and that manipulating such networks could reverse specific HFD-induced memory deficits.

We recently showed that object-based memory is impaired in HFD-exposed animals and that silencing of the ventral HPC (vHPC) with Designer Receptor Exclusively Activated by Designer Drugs (DREADD) rescued this memory deficits (Naneix et al., 2021). We therefore hypothesized that impairments in distinct components of object-based memory, namely recognition and location, in HFD animals could result from alterations of distinct vHPC efferent pathways. We focused on vHPC projections onto the nucleus accumbens (NAc) and the medial prefrontal cortex (mPFC), because they represent two of the main monosynaptic targets of the vHPC (Britt et al., 2012; Cenquizca and Swanson, 2007; Ciocchi et al., 2015; Gergues et al., 2020; Liu and Carter, 2018), they modulate object-based memory (Barker et al., 2017; Nelson et al., 2011; Sargolini et al., 2003) and we and others previously demonstrated that, in addition to the HPC, periadolescent HFD affects both NAc and mPFC functions (Ducrocq et al., 2019;

Labouesse et al., 2017, 2013; Naneix et al., 2017; Reichelt et al., 2019; Yaseen et al., 2019). We found that both pathways were hyperactive in HFD animals after object-based learning. Using an intersectional chemogenetic approach to manipulate selectively each pathway, we demonstrated specific beneficial effects of silencing either NAc- or mPFC-projecting vHPC neurons on HFD-induced deficits in either object recognition memory (ORM) or object location memory (OLM), respectively.

## Results

### Characterization of vHPC-NAc and vHPC-mPFC pathways

We first confirmed that vHPC-to-NAc and vHPC-to-mPFC pathways represent major vHPC efferent projections. Using Ai14(RCL-tdT)-D mice injected with an AAV-CaMKII-Cre in the vHPC (Supplementary Figure 1), numerous fibres expressing tdTomato were detected in the NAc (Figure 1A-B) and to a lesser extend in the mPFC (Figure 1C-D).

**Figure 1.**
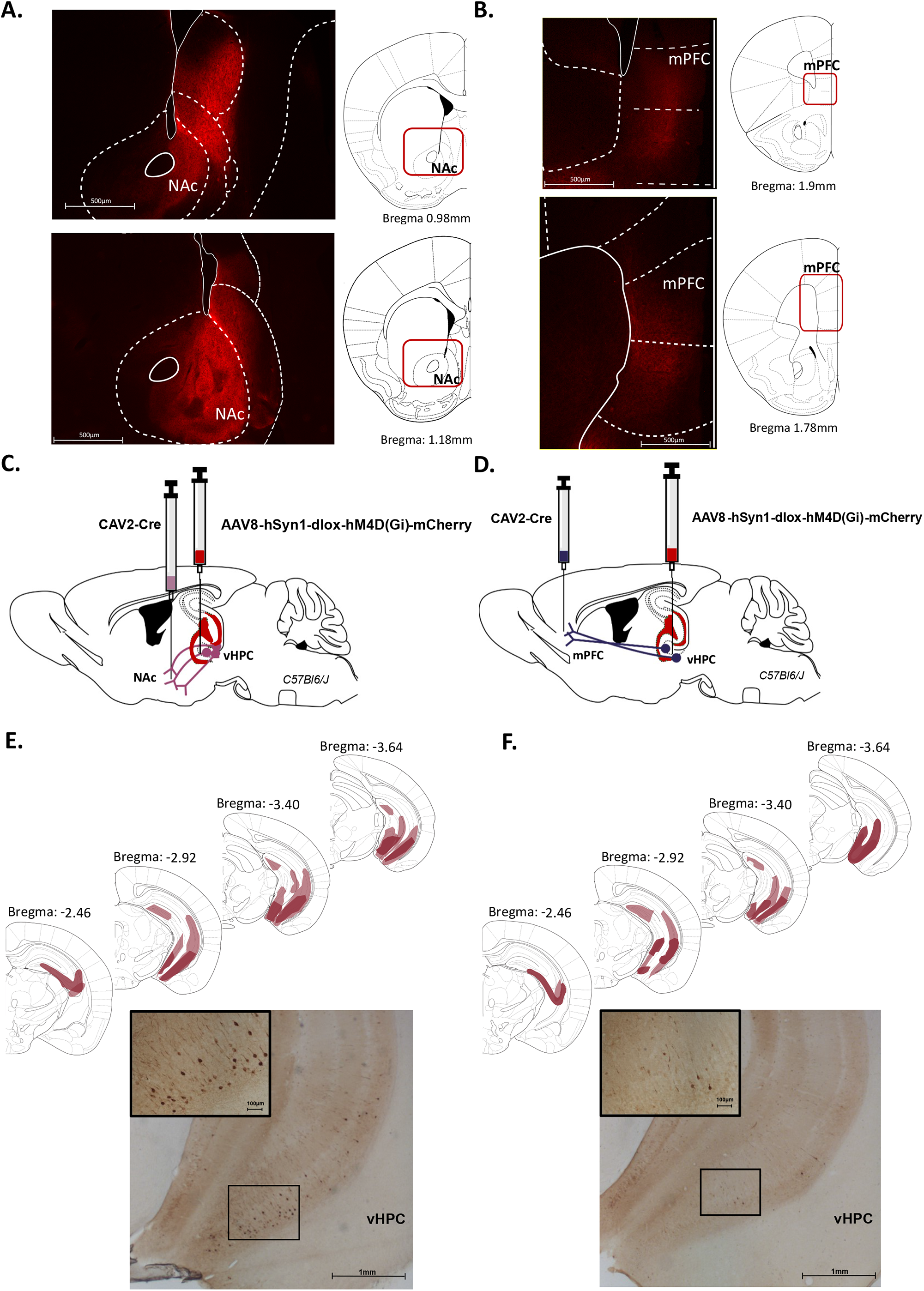
Characterization of ventral hippocampus projections to nucleus accumbens and medial prefrontal cortex. (A-B) Representative images illustrating expression of TdTomato in fibres in the nucleus accumbens (NAc, A) and the medial prefrontal cortex (mPFC, B) after AAV-CaMKII-Cre injection in the ventral hippocampus (vHPC) of Ai14(RCL-tdT)-D mice. (C-D) Schema of intersectional chemogenetic approach. An AAV-hSyn1-dlox-hM4D(Gi)-mCherry vector was injected into the vHPC, while a retrograde CAV2-Cre vector was injected in (E) the NAc or (F) the mPFC. Expression of mCherry is depicted for each condition after amplification using immunohistochemistry. Schematics adapted from (Paxinos and Franklin, 2004), indicating the largest (light red) or the smallest (dark red) viral infection. Representative images illustrating mCherry expression after CAV2-Cre injection in (E) the NAc or (F) the mPFC. Scale bar is set to 100μm, 500 μm or 1mm.

To specifically target vHPC-to-NAc and vHPC-to-mPFC pathways, we then used an intersectional strategy with a retrograde CAV2-Cre injected in either NAc or mPFC and Cre-responsive AAV-hM4Di-mCherry injected in vHPC. We detected strong mCherry labelling in the vHPC, particularly in the ventral CA1 and subiculum, after CAV2-Cre injection in the NAc demonstrating dense NAc-projecting vHPC neurons (Figure 1E) and a weaker vHPC labelling after CAV2-Cre infusion in the mPFC, revealing mPFC-projecting vHPC neurons in the ventral CA1 and subiculum (Figure 1F).

### Chemogenetic inhibition decreased HFD-induced activation of vHPC-NAc and vHPC-mPFC pathways

Mice of comparable body weight were randomly exposed to either CD (n= 15) or HFD (n= 61, 3 groups: vHPC-NAc n= 23, vHPC-mPFC n= 22 and control n= 16). After 12 weeks of diet exposure, fat mass and body weight differed between the groups [one-way ANOVA: F_(3,72)_= 8.6, p< 0.0001 and F_(3,72)_= 4.2, p< 0.01, respectively]. Post-hoc analysis showed a significant difference between CD and all HFD groups for fat content (p< 0.02, other comparisons: p> 0.26) and between CD and HFD vHPC-mPFC groups only for body weight (p< 0.01, other comparisons: p> 0.16; Table 1). These results confirm that periadolescent HFD increases fat accumulation in all groups compared to CD-fed mice.

**Table 1:** Body weight and fat mass measurements. Results are expressed as a percentage of fat mass versus body weight. *: p< 0.05, **: p< 0.01, ***: p< 0.001 compared to CD (one-way ANOVA followed by Tukey’s post-hoc test).

| Parameter/Group | CD<br>(n=15) | HFD mCherry<br>(n=16) | HFD vHPC-NAc<br>(n=23) | HFD vHPC-PFC<br>(n=22) |
| --- | --- | --- | --- | --- |
| **Body weight (grams)** | 30.0± 0.58 | 31.36± 0.61 | 31.5± 0.63 | 33.26± 0.67** |
| **% of fat content** | 6.66± 1.06 | 17.18± 2.89* | 19.77± 2.19*** | 23.27± 2.39*** |

As we previously reported a higher c-Fos expression in the dorsal hippocampus of HFD-fed mice after object-based training (Biyong et al., 2021), we evaluated whether similar HFD-induced overactivation is present in the vHPC and whether inhibiting NAc- and mPFC-projecting vHPC cells would prevent this activation. Therefore, c-Fos+ cells were quantified in CA1-subiculum, CA3 and Dentate Gyrus (DG) of the vHPC, after exposing animals to a novel context containing two identical novel objects (Figure 2A). CD and HFD groups similarly explored the novel objects excluding any behavioural influences on c-Fos expression between groups (Table 2). As similar levels of c-Fos+ cells were found between HFD control-CNO and HFD hM4Di-vehicle groups, indicating an absence of CNO effect by itself on c-Fos expression, they were merged into a single HFD control group (n= 13). One-way ANOVA revealed a significant difference between the 4 groups in the ventral CA1 [F_(3,30)_= 3.1, p= 0.04], with higher number of c-Fos+ cells in HFD control group than CD group (p= 0.05, other comparisons p> 0.17; Figure 2A-C), but not in the CA3 and DG [CA3: F_(3,30)_= 1.8, p= 0.16; DG: F_(3,30)_= 2.2, p= 0.10; Supplementary Figure 2].

**Table 2:** Object training before cellular imaging. No significant group difference was obtained.

| Parameter/Group | CD<br>(n=7) | HFD hM4Di<br>saline (n=7) | HFD mCherry<br>CNO (n=6) | HFD vHPC-NAc<br>CNO (n=7) | HFD vHPC-PFC<br>CNO (n=7) |
| --- | --- | --- | --- | --- | --- |
| **Time (s) object 1** | 20.7± 3.75 | 24.1± 2.74 | 23.12± 1.19 | 21.86± 2.29 | 22.09± 2.35 |
| **Time (s) object 2** | 18.39± 3.39 | 20.7± 0.92 | 22.24± 1.94 | 21.25± 2.36 | 21.36± 1.37 |
| **Total time (s) 1+2** | 39.73± 6.48 | 44.85± 3.57 | 45.36± 3.03 | 43.11± 4.22 | 43.45± 3.38 |
| **mCherry+ cells/mouse** | 158±48 | 157±16 | 204±33 | 155±19 | 120±30 |
| **Total mCherry+ cells** | 1109 | 942 | 1431 | 1088 | 839 |

**Figure 2.**
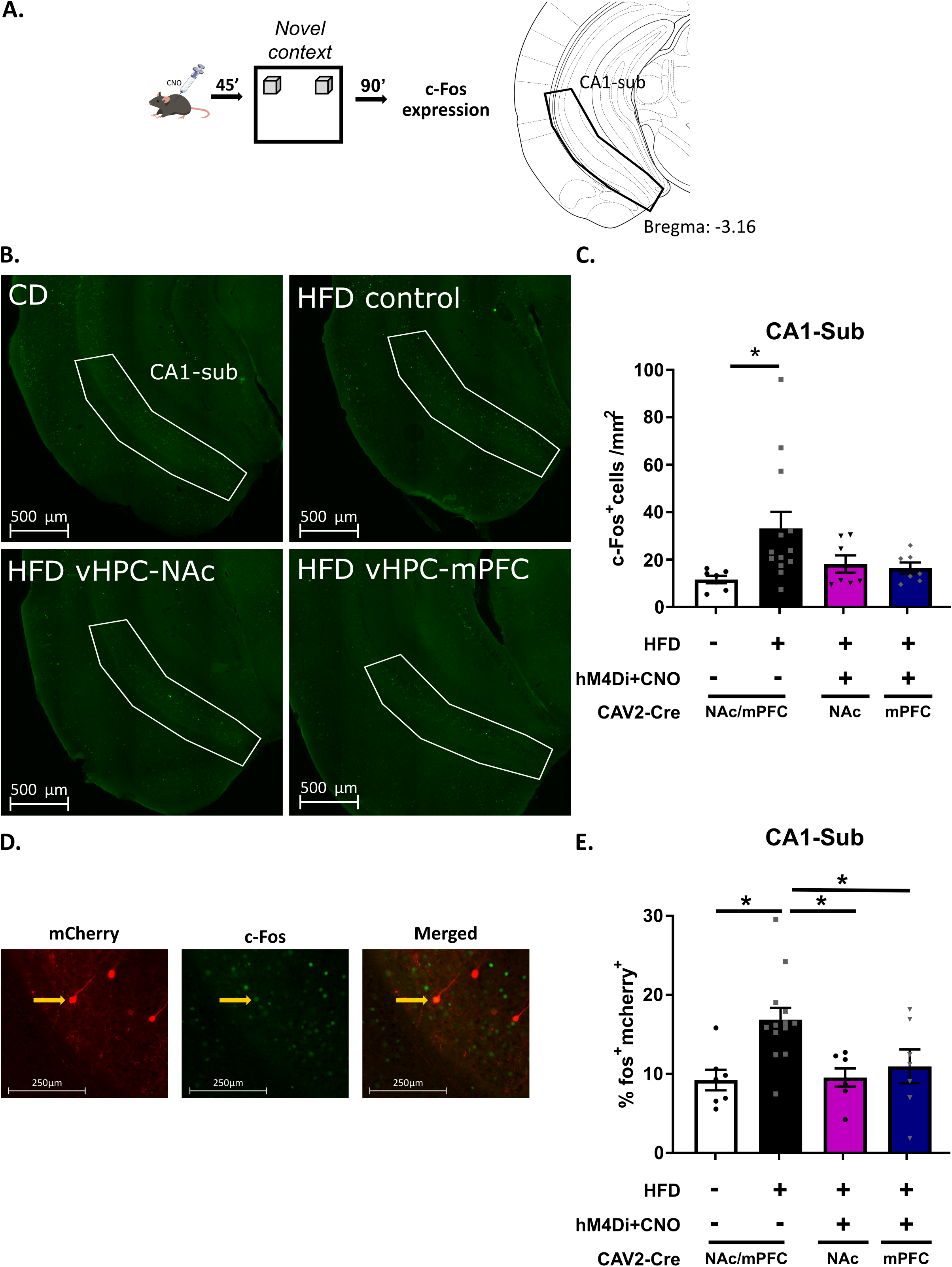
Effects of high-fat diet and chemogenetic silencing of vHPC-Nac or vHPC-mPFC pathway on c-Fos expression in ventral CA1/subiculum. (A) Schema of exposure to a novel arena containing two identical and novel objects. (B) Representative images of c-Fos-positive (Fos+) cells in the ventral CA1/subiculum for each group. (C) Quantification of c-Fos+ cells in the ventral CA1/subiculum of the different groups. Data are shown as the number of c-Fos+ cells per mm^2^. The area corresponding to ventral CA1/subiculum area is delineated. Scale bar is set at 500nm. (D) Representative example of mCherry and c-Fos labelling in the ventral CA1/subiculum. (E) Percentage of mCherry+ and c-Fos+ cells over total number of mCherry+ in the different groups. Scale bar is set to 250nm and data are shown as a percentage of cells. *: p<0.05 (one-way ANOVA followed by post-hoc test).

We then quantified the percentage of vHPC-to-NAc and vHPC-to-mPFC neurons (mCherry+) expressing c-Fos (Figure 2D-E; Table 2). Again, HFD control-CNO and HFD hM4Di-vehicle groups were merged in HFD control group (n= 13) as they displayed similar levels of mCherry labelled neurons double-stained for c-Fos. One-way ANOVA showed a significant difference among the 4 groups for the percentage of mCherry+/c-Fos+ cells [F_(3,30)_= 5.9, p= 0.002] with HFD control group presenting significantly higher proportion of double labelled cells than the 3 other groups (p= 0.009 vs CD, p= 0.01 vs HFD vHPC-NAc and p=0.05 vs vHPC-mPFC), that did not differ from each other (p> 0.9; Figure 2E).

Altogether, these results show that 1) HFD enhanced c-Fos activation in the ventral CA1-subiculum and more specifically in NAc- and mPFC-projecting vHPC neurons and 2) silencing of either vHPC pathway normalized HFD-induced over-activation, therefore validating our intersectional approach.

### Silencing of vHPC-NAc, but not vHPC-mPFC, pathway rescued HFD-induced long-term ORM deficits

As chemogenetic vHPC silencing during ORM training restored HFD-induced long-term memory deficits (Naneix et al., 2021), we wondered to which extent the specific silencing of either NAc- or PFC-projecting vHPC cells upon CNO injection 45min before training could rescue these deficits (Figure 3A). Four groups were used for this purpose: CD, HFD control, HFD vHPC-NAc and HFD vHPC-mPFC (8-12 mice/group; see Table 3 for objects exploration during training and test). During the ORM test performed 24 hours later, one-way ANOVA revealed a significant difference among the 4 groups [F_(3,35)_= 4.5, p= 0.009] with the percentage of preference for the novel object being significantly higher in CD and HFD vHPC-NAc groups than in HFD control (p< 0.02). No other group difference was observed (p> 0.2). Indeed, CD-fed mice explored preferentially the novel object [one sample t-test against 50%: t_(7)_=6.4, p= 0.0004] whereas HFD control mice exhibited ORM deficits [t_(7)_= 0.1, p= 0.92; Figure 3B]. Inactivation of the vHPC-NAc pathway during training restored HFD-induced ORM deficits [t_(11)_= 6.7, p< 0.0001], while vHPC-mPFC inactivation did not [t_(10)_= 1.6, p= 0.13; Figure 3B; Table 3]. These results indicate that periadolescent HFD induces long-term ORM deficits that are restored specifically by inactivating the vHPC-NAc pathway.

**Table 3:** Object recognition memory task. % expl. ratio represents a ratio of time exploring one object on total exploration time of both objects. *: p< 0.05 compared to CD (one-way ANOVA followed by Tukey’s post-hoc test).

| Parameter/Group | CD<br>(n=8) | HFD mCherry<br>(n=8) | HFD vHPC-NAc<br>(n=12) | HFD vHPC-PFC<br>(n=11) |
| --- | --- | --- | --- | --- |
| **ORM % expl. ratio** | 52.05± 2.01 | 52.96± 1.79 | 48.96± 2.71 | 50.85± 2.24 |
| **Time (s) object to be changed** | 10.41± 0.4 | 10.59± 0.36 | 9.79± 0.54 | 10.17± 0.45 |
| **Time (s) object not to be changed** | 9.59± 0.4 | 9.41± 0.36 | 10.21± 0.54 | 9.83± 0.45 |
| **Time (s) to reach criterium (training)** | 388± 32.89 | 274.4± 32.96 | 298.5± 32.41 | 301.4± 34.73 |
| **Time (s) to reach criterium (test)** | 370.1± 44.21 | 264± 33.51 | 229.1± 20.62* | 285.7± 26.14 |

**Figure 3.**
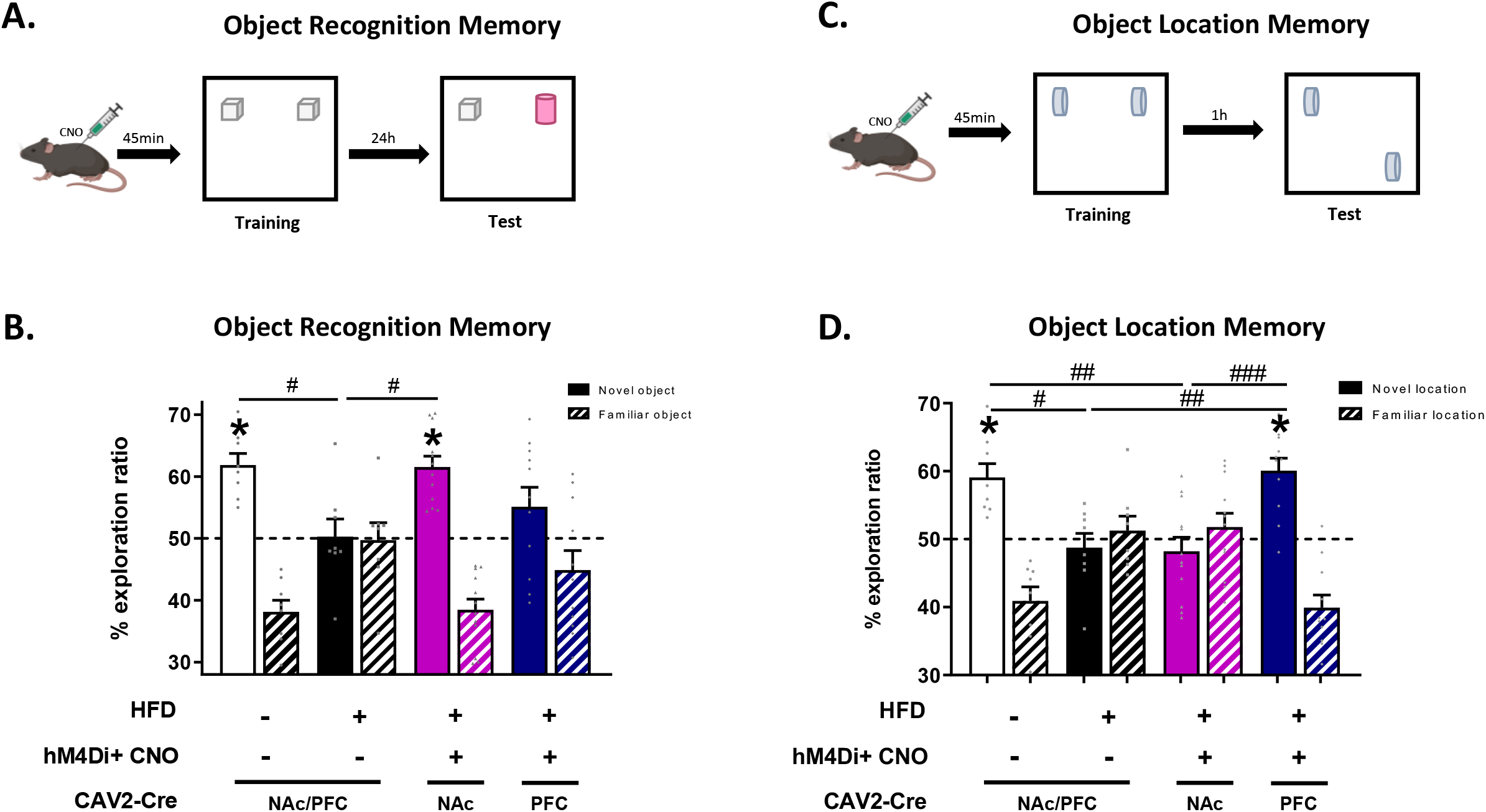
Impacts of high-fat diet and chemogenetic silencing of vHPC-Nac or vHPC-mPFC pathway on object-based memory. (A) Schema of object recognition memory (ORM) task. (B) ORM performance expressed as percentage of exploration time of novel (empty bars) or familiar (striped bars) object over both objects. (C) Schema of object location memory (OLM) task. (D) OLM performance expressed as percentage of exploration time of novel (empty bars) or familiar (striped bars) location over both objects. *: p<0.05 (one-sample t-test, different from 50%); #: p<0.05, ###: p<0.001 (one-way ANOVA followed by post-hoc test).

### Silencing of vHPC-mPFC, but not vHPC-NAc, pathway rescued HFD-induced OLM deficits

We also evaluated the effect of specifically silencing the vHPC pathways on another object-based memory task, the object location memory (OLM), known to be impaired by periadolescent HFD (Glushchak et al., 2021; Khazen et al., 2019; Valladolid-Acebes et al., 2013). As for ORM, 4 groups (CD, HFD control, HFD vHPC-NAc and HFD vHPC-mPFC; 8-12 mice/group) were injected with CNO 45min before training (Figure 3C; Table 4). During the OLM test performed 1 hour later, a significant difference was found among the 4 groups for the percentage of preference for the displaced object [F_(3,35)_= 10.11, p<0.0001], with the CD and HFD vHPC-mPFC groups showing a significantly higher preference than HFD control and HFD vHPC-NAc (p< 0.01). No other group difference was observed (p> 0.9). CD-fed mice explored preferentially the displaced object [one sample t-test against 50%: t_(7)_= 4.4, p=0.003] whereas HFD control mice exhibited ORM deficits [t_(7)_= 0.59, p= 0.57; Figure 3D]. Chemogenetic silencing of the vHPC-NAc pathway before training failed to improve OLM performance [t_(11)_= 0.9, p= 0.41] whereas manipulation of the vHPC-mPFC pathway improved OLM performance in HFD animals [t_(10)_= 5.5, p<0.001; Table 4]. These results indicate that pre-training or pre-test inactivation of the vHPC-mPFC pathway, but not the vHPC-NAc, restored HFD-induced OLM deficits.

**Table 4:** Object location memory task. % expl. ratio represents a ratio of time exploring one object on total exploration time of both objects. No significant group difference was obtained.

| Parameter/Group | CD<br>(n=8) | HFD mCherry<br>(n=8) | HFD vHPC-NAc<br>(n=12) | HFD vHPC-PFC<br>(n=11) |
| --- | --- | --- | --- | --- |
| **OLM % expl. ratio** | 46.64± 1.55 | 48.88± 1.53 | 50.64± 1.49 | 47.71± 1.86 |
| **Time (s) object to be displaced** | 9.33± 0.31 | 9.77± 0.31 | 10.13± 0.3 | 9.54± 0.37 |
| **Time (s) object not to be displaced** | 10.67± 0.31 | 10.22± 0.3 | 9.873± 0.3 | 10.46± 0.37 |
| **Time (s) to reach criterium (training)** | 318.1± 50.59 | 330.1± 33.27 | 340.8 ±37.28 | 358.4 ±24.74 |
| **Time (s) to reach criterium (test)** | 347± 42.05 | 303.8± 46.88 | 337.9 ± 27.8 | 339.6 ±36.99 |

### HFD and projecting vHPC cells manipulation did not affect anxiety-like behaviour

HFD intake (Fulton et al., 2022) or manipulation of specific vHPC pathways (Padilla-Coreano et al., 2016; Parfitt et al., 2017) has been reported to affect anxiety-like behaviour. We therefore evaluated, using the EPM test, whether anxiety was affected in our conditions. One-way ANOVA revealed no difference between the 4 groups (8-11 mice/group) regarding the ratio of time spent in open arms [F_(3,33)_ = 1.4, p= 0.26; Supplementary Figure 3, left] or the number of entries [F_(3,33)_= 0.33, p= 0.8; Supplementary Figure 3, right]. These results indicate that neither periadolescent HFD nor vHPC pathway manipulation affect anxiety-like behaviour, and therefore exclude that any effects on memory performance would be due to differences in anxiety levels among the groups.

## Discussion

The current study provides evidence that periadolescent HFD induced deficits in non-aversive and non-rewarded object-based memory in mice, without affecting anxiety-like behaviour. These memory deficits were associated with enhanced activation of vHPC neurons projecting to NAc and mPFC. Using a pathway specific chemogenetic approach that was effective in normalising this activation, we demonstrated that 1) silencing of NAc-projecting, but not mPFC-projecting, vHPC neurons restored long-term ORM deficits induced by HFD whereas 2) inactivation of mPFC-projecting, but not NAc-projecting, vHPC neurons rescued HFD-induced OLM deficits. Our results therefore revealed a double dissociation in the beneficial outcomes of vHPC-NAc or vHPC-mPFC silencing on HFD-induced ORM and OLM deficits, without any effect on anxiety-like behaviour.

Our previous studies revealed that 12 weeks of periadolescent HFD induced a higher c-Fos activation of the dorsal hippocampus after object-based training (Biyong et al., 2021). Here, we obtained similar c-Fos overexpression in the CA1/subiculum of the vHPC after object training, as well as in projecting areas NAc shell and mPFC (data not shown), providing evidence that a large neural network is overactive in HFD-fed mice following object training. In HPC, HFD effect is more pronounced in ventral CA1 and subiculum, which represent the main output of the vHPC (Britt et al., 2012; Cenquizca and Swanson, 2007; Ciocchi et al., 2015; Gergues et al., 2020; Liu and Carter, 2018). This could be related to HFD-induced morphological and electrophysiological changes in CA1 as it was reported that 8-12 weeks of post-weaning HFD intake enhances dendritic spine density in CA1 pyramidal neurons (Valladolid-Acebes et al., 2013) and induces aberrant *in vivo* long-term potentiation in CA1 (Vouimba et al., 2021). Thanks to our intersectional virus approach, we were able to restrict our c-Fos analyses to NAc-projecting or mPFC-projecting vHPC neurons located in ventral CA1/subiculum. Periadolescent HFD induced an increased percentage of c-Fos expressing neurons in both pathways after object training, corroborating what was found at the level of the whole ventral CA1-subiculum. Importantly, CNO decreased the activation of vHPC-to-NAc and vHPC-to-mPFC neurons expressing inhibitory DREADDs in HFD-fed mice, validating the efficacy of our intersectional chemogenetic approach. This restricted chemogenetic inhibition tends to attenuate HFD-induced overactivation of the whole ventral CA1/subiculum, possibly through a network effect.

Our results indicate that periadolescent HFD induces long-term ORM deficits, corroborating our recent studies showing specific HFD effects on long-term, but not short-term, ORM (Biyong et al., 2021; Naneix et al., 2021). We had previously reported that chemogenetic inactivation of vHPC projecting neurons during training, by restricting DREADDs expression primarily to excitatory principal neurons thanks to the use of a CaMKII promoter, is able to restore HFD-induced ORM deficits^27^. Here, we obtained similar ORM improvement in HFD-fed animals using specific inactivation of NAc-projecting vHPC neurons during memory formation. This finding is in accordance with previous results demonstrating the role of NAc in ORM consolidation (Sargolini et al., 2003) and in integrating hippocampal-derived information during memory consolidation (Kerfoot and Williams, 2018; Roozendaal et al., 2001). Our results also support previous findings indicating that HFD affects NAc functions after periadolescent HFD exposure (Ducrocq et al., 2019; Labouesse et al., 2013; Naneix et al., 2017). Finally, chemogenetic silencing of mPFC-projecting vHPC neurons did not rescue HFD-induced ORM deficits corroborating recent results reporting that manipulation of vHPC-mPFC pathway did not affect ORM in control animals (Naneix et al., 2017; Phillips et al., 2019).

Previous findings indicate that periadolescent exposure to HFD drastically affects spatial memory assessed in the Morris water maze or the radial arm maze (Boitard et al., 2014, 2012) but also using non-aversive and non-rewarded OLM (Glushchak et al., 2021; Khazen et al., 2019; Valladolid-Acebes et al., 2013). We here demonstrate that silencing of the vHPC-mPFC, but not the vHPC-NAc, pathway restores OLM deficits induced by periadolescent HFD. Our results are related to previous studies indicating that periadolescent HFD altered mPFC functions (Labouesse et al., 2017; Reichelt et al., 2019; Yaseen et al., 2019) and that mPFC lesion or inhibition of vHPC-mPFC pathway in control animals disrupted spatial memory performance but left ORM intact (Barker et al., 2017; Morici et al., 2022; Nelson et al., 2011). It must be stressed that OLM was tested one hour after training. As CNO action lasts several hours (Alexander et al., 2009), mPFC-vHPC pathways was inactivated during both OLM training and test. Additional experiments will determine whether chemogenetic inactivation of vHPC-mPFC pathways after training and before test still has a beneficial effect on OLM performance therefore supporting mPFC role in orchestrating action to perform a given task (Friedman and Robbins, 2022).

Recent studies indicate that a very small proportion (5-10%) of vHPC neurons project to two or three areas (Ciocchi et al., 2015; Gergues et al., 2020; Parfitt et al., 2017). However, the double dissociation in our effects of pathway-specific silencing on ORM and OLM precludes that collaterals of mPFC-projecting and NAc-projecting vHPC neurons may be involved in our effects. Also, the double dissociation observed in the present study excludes any influences of sensory, motor, motivational or attentional processes, which would have similarly affected both memory tasks. Finally, our diet or pathway manipulation did not affect anxiety-like behaviour, ruling out that our effects on memory were due to differences in anxiety levels among the groups. This is in line with previous studies performed after juvenile or periadolescent HFD exposure (Boitard et al., 2015; Khazen et al., 2019) but in contrast with others showing that adult HFD exposure triggered anxiety-like behaviour (Décarie-Spain et al., 2018; Fulton et al., 2022). This suggests a developmental HFD impact on anxiety which deserves more investigation.

## Conclusions

The current study provides evidence that periadolescent HFD induces deficits in different types of memory through an over-activation of specific vHPC efferent pathways and dampening this activation alleviates these memory deficits. These findings extend our knowledge about the cerebral impact of obesogenic diet focusing on brain network and connectivity and emphasize the distinct role of specific vHPC efferent pathways in different types of object-based memory. Future studies are required to examine the effects of manipulation of vHPC pathways, including those to anterior olfactory nucleus (Aqrabawi and Kim, 2018), in other types of memory impairments promoted by HFD, such as social and olfactory memory (Reichelt et al., 2019; Yaseen et al., 2019). Moreover, further investigation is also necessary to decipher what is the origin of this hippocampal over-activation in obesogenic diet-fed rodents.

According to the important role played by HPC hyperactivity in memory decline associated with normal and pathological aging (Bakker et al., 2012; Yassa et al., 2011) as well as disruption of vHPC efferent pathways in neuropsychiatric disorders (Bagot et al., 2015; Li et al., 2015; Phillips et al., 2019), future research has to contribute to the development of novel therapeutic strategies to alleviate disorders characterised by impaired hippocampal homeostasis.

## Materials and Methods

### Animals, diet and housing

Male C57BL/6 J mice aged of 3 weeks old (Janvier Labs, France) were divided randomly into groups of 8 per cage (45x 25x 20 cm, containing a paper house, nesting material and a small wooden stick) and had *ad libitum* access to a control diet (CD; n=16; 2.9 kcal/g; 8% lipids, 19% proteins, 73% carbohydrates mostly from starch; A04, SAFE) or a high-fat diet (HFD; n=64; 4.7 kcal/g; 45% lipids mostly saturated fat from lard, 20% proteins, 35% carbohydrates mainly from sucrose; D12451, Research Diet). We focused on males given that female rodents do not consistently exhibit memory deficits after post-weaning obesogenic diet intake (Abbott et al., 2016; Hwang et al., 2010), which is probably related to ovarian hormones that appear to protect females from obesity and metabolic impairments (Palmer and Clegg, 2015). All animals were housed in a temperature-controlled room (22 ±1°C) maintained under a 12 h light/dark cycle (lights on at 8:00 am, lights off at 8:00 pm) and had free access to food and water during 12 weeks (before the beginning of behaviour) as well as during all behavioural procedures before euthanasia (13-14 weeks of diet exposure in total). Animals were weighed at arrival, before and after surgery, before each behavioural test as well as before euthanasia. All animal care and experimental procedures were in accordance with the INRAE Quality Reference System and with both French (Directive 87/148, Ministère de l’Agriculture et de la Pêche) and European legislations (Directive 86/609/EEC). They followed ethical protocols approved by the Region Aquitaine Veterinary Services (Direction Départementale de la Protection des Animaux, approval ID: B33-063-920) and by the animal ethic committee of Bordeaux CEEA50. Every effort was made to minimize suffering and reduce the number of animals used. Both surgeries and behavioural experiments were performed at adulthood. The day before euthanasia, fat mass (in grams) was measured by nuclear magnetic resonance (minispec LF90 II, Bruker, Wissembourg, 67166) (Naneix et al., 2021) and divided by body weight (in grams) as a ratio for each mouse. Three homozygous Ai14 Cre reporter adult mice [B6.Cg-Gt(ROSA)^26Sortm14(CAG-tdTomato)Hze^/J or simply Ai14(RCL-tdT)-D; Jackson laboratory] under CD were also used to characterize vHPC-to-Nac and vHPC-to-mPFC pathways allowing the Cre-dependent expression of the red fluorophore td-Tomato specifically in vHPC projecting neurons and their efferents (see below).

### Viral vectors and drugs

An Adeno Associated Virus (AAV) carrying the Cre recombinase was used for the experiment performed in the Ai14(RCL-tdT)-D mice (AAV1-CaMKII-Cre, 4.6 ×10^12^ vg/ml; ETH VVF, Zurich, RRID v315-1). For the chemogenetic manipulation of specific pathways, an anterograde AAV carrying Cre-dependent inhibitory DREADD (AAV8-hSyn1-dlox-hM4DGi-mCherry, 5.4 ×10^12^ vg/ml; ETH VVF, Zurich, RRID v84-8) or control virus (AAV8-hSyn1-dlox-mCherry, 6.3 ×10^12^ vg/ml; ETH VVF, Zurich, RRID v116-8) were used in combination with a retrograde canine virus (CAV2) carrying the Cre recombinase (CAV2-Cre). This led to 4 groups: 2 control groups (CD and HFD control) injected with AAV control and CAV2-Cre and 2 experimental groups (HFD vHPC-NAc and HFD vHPC-mPFC) injected with AAV-dlox-hM4Di and CAV2-Cre.

The exogenous DREADD ligand Clozapine-N-Oxide (CNO, Enzo Life Sciences, RRID BML-NS105) was dissolved in 0.9% saline for a final concentration of 2 mg/kg. Saline solution was used for vehicle injections. Both CNO and vehicle were freshly prepared every day and delivered by intraperitoneal injection (10ml/kg) 45min before each behavioural test.

### Stereotaxic surgery

After 7–8 weeks under CD or HFD, mice were anaesthetized under isoflurane (5% induction; 1–2% maintenance), injected with the analgesic buprenorphine (Buprecar, 0.05 mg/kg s.c.) and the non-steroidal anti-inflammatory drug carproxifen (Carprox, 5mg/kg s.c.) and were placed on a stereotaxic apparatus (David Kopf Instruments). The scalp was shaved, cleaned and locally anaesthetized with a local subcutaneous injection of lidocaine (Lurocaine, 0.1ml). Viral vectors were infused using a 10 μL Hamilton syringe (Hamilton) and an ultra-micro pump (UMP3, World Precision Instruments, USA). For the experiment performed in the Ai14(RCL-tdT)-D mice, two injections per side (AP −3.2 mm, ML ± 3.2 mm from Bregma, DV −3 and −4mm from the skull surface, according to ^59^) of 1 μL each were performed with the vector AAV1-CaMKII-Cre. For the dual virus experiments, 1 μL of the AAV (AAV-dlox-hM4Di or AAV control) was injected over 6 min (150 nL/min) in the vHPC at 1 site in each hemisphere, coordinates were AP −3.2 mm, ML ± 3.2 mm from Bregma, DV −4mm from the skull surface. Then 250nl (mPFC; AP +1.9 mm, ML ± 0.3 mm, DV −3 mm from skull) or 400nl (NAc shell; AP +1.2 mm, ML ± 0.6 mm, DV −4.8 mm from skull) of CAV2 virus were injected at a rate of 100 nL/min over 2min 30sec and 4min, respectively. In all cases, the pipette was left in place for a 5 minutes diffusion period, before being slowly removed. The incision was closed with sutures and the animal was kept on a heating pad until recovery. Mice were single housed for 4 days and their body weight and behaviour were closely monitored. Then, they were housed in groups of 4 mice per cage and 4 weeks later (allowing optimal virus expression) behavioural tests started.

### Behavioural procedures

The 4 groups (CD, HFD control, HFD vHPC-NAc and HFD vHPC-mPFC; 8-12 mice/group) were submitted to different object-based memory tasks known to be impaired by periadolescent HFD, namely long-term object recognition memory (ORM) (Biyong et al., 2021; Naneix et al., 2021) and short-term object location memory (OLM) (Glushchak et al., 2021; Khazen et al., 2019; Valladolid-Acebes et al., 2013). Anxiety-like behaviours were also evaluated in an elevated plus-maze (EPM). The order of tests was performed in a random way. Animals were handled twice a day for three days before the beginning of the first behavioural test. All behavioural tests were performed during light-phase and under white light (15 lux). Mice of the different groups were tested in a pseudo-random order by an experimenter blind to group identity. Between each trial, arena (and objects where appropriate) were cleaned with 10% of alcohol. All behaviour analysis was scored online, apart from the EPM that was automatically analysed by SMART system (Bioseb, Vitrolles, France).

### Object Recognition Memory (ORM)

During training, two identical new objects were presented in a novel open field arena (40x 40x 40cm, wood) and each mouse was left freely to explore them. Long-term memory was assessed twenty-four hours later, by randomly replacing one of the objects by a novel one. In both phases, object exploration was defined as nose and whiskers pointed towards the object in a distance of less than 1-1.5cm away and an inclusion criterion of 20 seconds of total exploration was set at a 10 minutes exploration maximum (Leger et al., 2013), otherwise mice were excluded from analysis. Data were calculated as the % of exploration ratio of novel (or familiar) versus total object exploration time.

### Object Location Memory (OLM)

Two identical new objects (different from ORM) were presented during training in a new room with a new open field arena (40x 40x 40cm, plastic) and each mouse was left freely to explore them. Short-term memory was assessed one hour later, by randomly placing one of the objects in a novel location. In both phases, an inclusion criterion of 20 seconds of total exploration was set at a 10 minutes exploration maximum, otherwise mice were excluded from analysis. Data were calculated as the % of exploration ratio of novel (or familiar) location versus total object exploration time.

### Elevated Plus-Maze (EPM)

Mice were allowed to freely explore the plus-shaped acrylic maze sized 30 × 8 × 15 cm (closed arms) and 30 × 8 cm (open arms) connected by a central part (8 × 8 cm) for 10 min. The maze is elevated 120 cm above the floor. A mouse was considered to enter one zone only when it placed all four limbs in any particular part of the maze. Time spent in the open versus the closed arms was recorded and results are depicted as a ratio of time spent in open arms over total test time in seconds. In addition, a ratio of the number of entries in the open arms over total number of entries was calculated. An increased ratio for the open arms (time and/or number of entries) indicates low anxiety.

### Tissue collection and Immunochistochemistry

All mice were sacrificed after behavioral testing. For experiments evaluating c-Fos activation, some mice were sacrificed 90min after being placed in new open field arena (40x 40x 40cm) with two identical new objects (corresponding to ORM and OLM training) during 8 minutes. These mice were divided into 5 groups (of 6-7 mice each, 3 mice presenting very low number of mCherry labelled cells (< 30) in vHPC were excluded), 4 groups being injected with CNO 45min before the task (CD, HFD control-CNO, HFD vHPC-NAc and HFD vHPC-mPFC) and an additional HFD hM4Di-vehicle group (composed of 4 vHPC-NAc and 4 vHPC-mPFC mice) injected with vehicle to control for CNO effect.

Animals were deeply anaesthetized with a mix of pentobarbital and lidocaine (Exagon, 300mg/kg and Lurocaine, 30mg/kg) before being transcardiacally perfused with phosphate-buffered saline (PBS) solution followed by 4% paraformaldehyde (Sigma-Aldrich). Brains were collected, post-fixed in 4% paraformaldehyde at 4°C for 2 days, then switched to PBS solution and stored at 4°C until slicing. 40-μm coronal sections were cut via vibratome (Leica) and stored in cryoprotective solution (glycerol and ethylene glycol Sigma-Aldrich in PBS) at −20°C.

### DREADD expression

On the first day, slices were washed with PBS and incubated with 0.33% H_2_O_2_ solution in PBS for 30min. Then, they were incubated with blocking solution [0.2% Triton, 3% foetal bovine serum (FBS), in PBS] for 90min followed by an incubation with a rabbit anti-DsRed antibody (1:2000, in blocking solution; Takara Bio-Living Colors, RRID #632496) overnight at 4°C. Next, slices were incubated with a biotinylated goat anti-rabbit antibody (1:1000in PBS with 1% FBS; Vector Laboratories, RRID BA-1000) for 90min at room temperature, followed by an 1h incubation in avidin–biotin–peroxidase solution (Vectastain, Vector Laboratories, RRID PK-6200). Slices were then washed with PBS and Tris buffer solution (TBS, pH= 7.4) followed by diaminobenzidine (DAB, Sigma-Aldrich, RRID D5905) incubation for 15-30min (1 tablet of DAB and 1 tablet H_2_O_2_; 5ml of distilled H_2_O in 20ml of TBS). After stopping the reaction, slices were stored at 4°C. Slices were then mounted on gelatin-coated slides, covered by medium (Southern Biotech) and cover-slipped. Each section was photographed (Nikon-ACT-1 software).

### Pathway activation

A double immunofluorescence was performed. Slices were washed with PBS solution, incubated with blocking solution for 90min then with a combination of two primary antibodies: rabbit anti-Fos (1:2000; Cell Signaling Technology, RRID #2250) and chicken anti-mCherry (1:5000; Abcam, RRID ab150080), all in blocking solution of 3% FBS, 0.2% Triton in PBS (72h, at 4°C). Then, slices were incubated with a combination of two secondary antibodies, goat anti-chicken (1:1000, A488; Abcam, RRID ab205402) and goat anti-rabbit (1:1000, A594; Abcam, RRID ab150173). Slices were stored at 4°C until mounting on non-gelatin-coated slides followed by cover of DAPI fluoromount (Southern Biotech) and cover-slipped. All slices were photographed with Nanozoomer slide scanner Hamamatsu NANOZOOMER 2.0HT (Bordeaux Imaging Center, Univeristy of Bordeaux, France). QuPath program [QuPath v.0.3.0 (Bankhead et al., 2017)] was used for quantification of c-Fos positive cells.

## Materials availability statement

Weight, fat mass, immunohistochemical and behavioural data are available in Supplementary file.

## Statistics

Two mice died after surgery (one CD and one HFD vHPC-mPFC) and 2 mice were excluded after histological control (one HFD vHPC-mPFC and one HFD vHPC-NAc with unilateral labelling). Data were analyzed with Prism Software (GraphPad) and are expressed as mean ± SEM. Significance was set at p ≤ 0.05. For each group, ORM and OLM performance was compared against 50% exploration ratio (chance level) using one-sample *t*-test. One-way ANOVA followed by Tukey’s post hoc analysis was used to compare the 4 groups.

## Acknowledgements

The authors thank Gregory Artaxet and Eva Bruchet (NutriNeuro lab) for taking daily care of the animals, Léa Décarie-Spain for help during surgery, Lola Fauré for some illustrations and the Bordeaux Imaging Center (a service unit of University of Bordeaux and a national infrastructure, France BioImaging) where part of microscopy was completed. This work was supported by INRAE (to G.F.), CNRS (to E.C.), and French National Research Agency (ANR-14-CE13-0014 GOAL and ANR-15-CE17-0013 OBETEEN to E.C. and G.F. and ANR-16-CE37-0010 ORUPS to G.F.). I.B. was the recipient of a PhD fellowship from the French Ministry of Research and Higher Education (2018-2021).

## SUPPLEMENTARY INFORMATION

**Figure S1.**
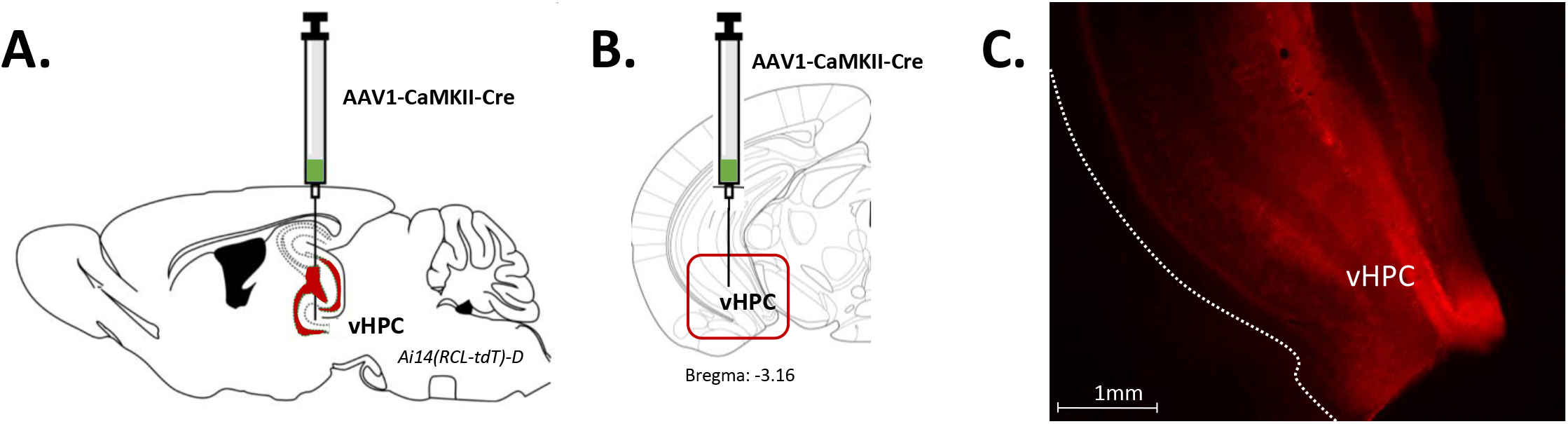
Cre injection in the ventral hippocampus of Td-Tomato mice. Related to Figure 1. Parasagittal (A) and frontal (B) schema illustrating AAV-CaMKII-Cre injection in the ventral hippocampus (vHPC) of Ai14(RCL-tdT)-D mice. (C), Representative image illustrating expression of TdTomato in vHPC neurons.

**Figure S2.**
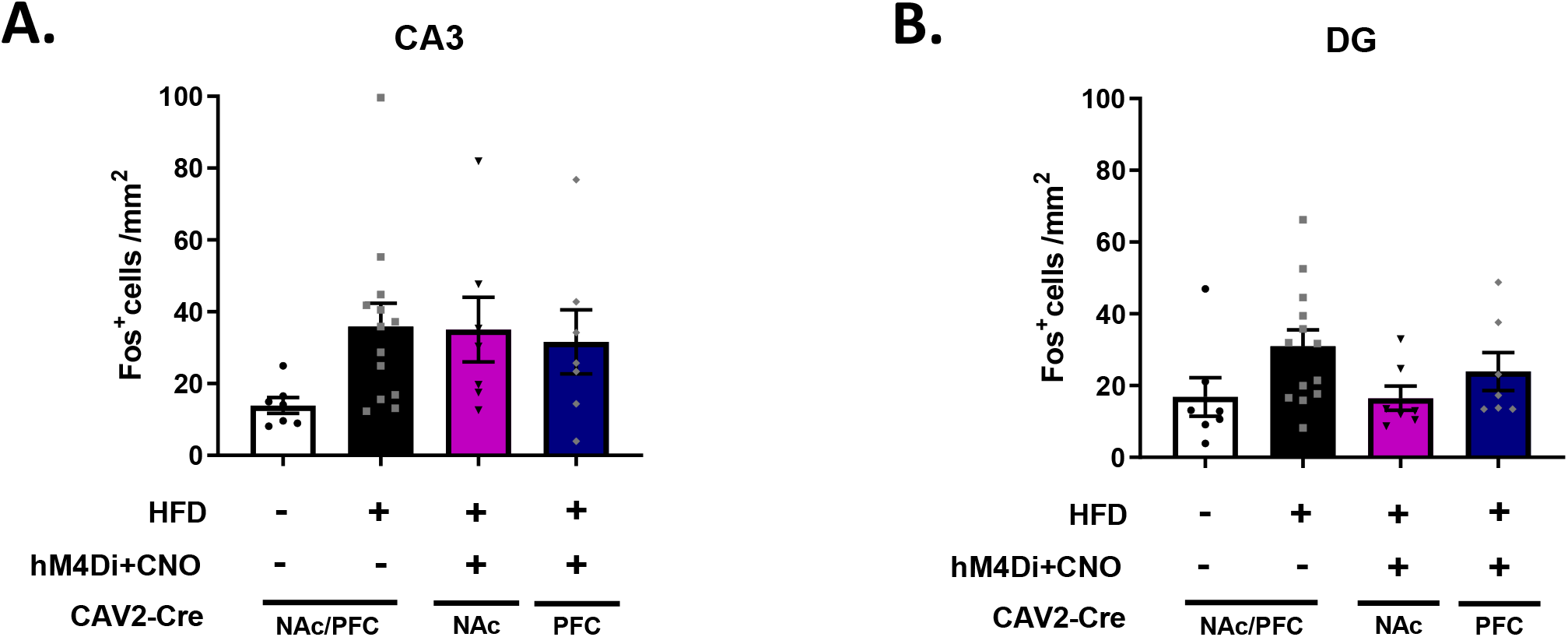
Effect of high-fat diet and chemogenetic manipulation on c-Fos expression in ventral CA3 and dentate gyrus. Related to Figure 2. Quantification of c-Fos-positive cells in the ventral CA3 (A) and dentate gyrus (B) of the different groups. Data are shown as the number of c-Fos-positive cells per mm^2^.

**Figure S3.**
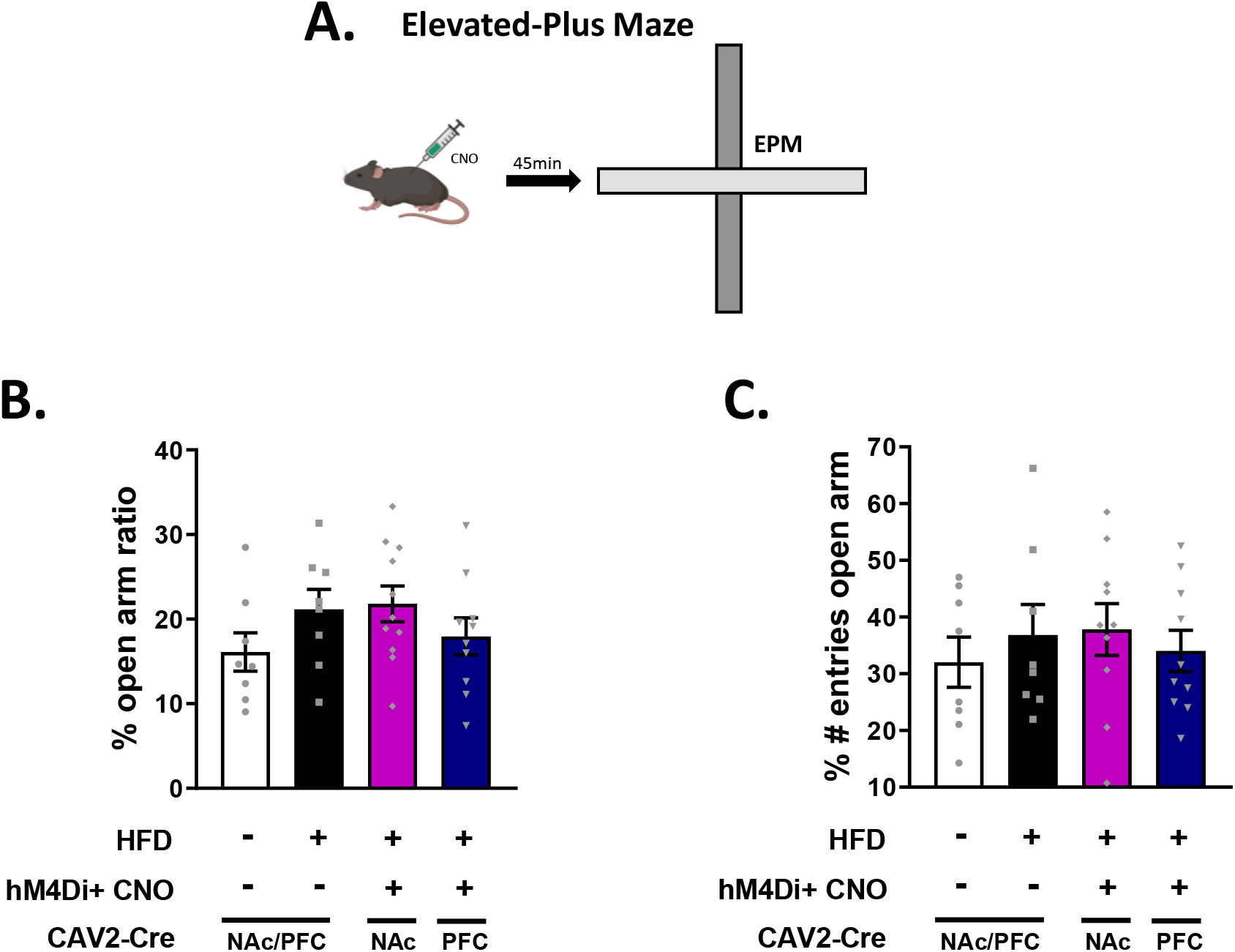
Effect of high-fat diet and chemogenetic manipulation on anxiety-like behaviors. Related to Figure 3. (A), Schema of elevated plus-maze task. Results presented as a percentage of time spent in open arms over total exploration time of both open and closed arms (B) and number of entries in open arms over total number of entries in both open and closed arms (C).

## References

Abbott KN, Morris MJ, Westbrook RF, Reichelt AC. 2016. Sex-specific effects of daily exposure to sucrose on spatial memory performance in male and female rats, and implications for estrous cycle stage. Physiol Behav 162:52–60. doi:10.1016/j.physbeh.2016.01.036

Alexander GM, Rogan SC, Abbas AI, Armbruster BN, Pei Y, Allen JA, Nonneman RJ, Hartmann J, Moy SS, Nicolelis MA, McNamara JO, Roth BL. 2009. Remote control of neuronal activity in transgenic mice expressing evolved G protein-coupled receptors. Neuron 63:27–39. doi:10.1016/j.neuron.2009.06.014

Andersen SL. 2003. Trajectories of brain development: point of vulnerability or window of opportunity? Neurosci Biobehav Rev 27:3–18. doi:10.1016/s0149-7634(03)00005-8

Aqrabawi AJ, Kim JC. 2018. Hippocampal projections to the anterior olfactory nucleus differentially convey spatiotemporal information during episodic odour memory. Nat Commun 9:2735. doi:10.1038/s41467-018-05131-6

Bagot RC, Parise EM, Peña CJ, Zhang H-X, Maze I, Chaudhury D, Persaud B, Cachope R, Bolaños-Guzmán CA, Cheer JF, Cheer J, Deisseroth K, Han M-H, Nestler EJ. 2015. Ventral hippocampal afferents to the nucleus accumbens regulate susceptibility to depression. Nat Commun 6:7062. doi:10.1038/ncomms8062

Bakker A, Krauss GL, Albert MS, Speck CL, Jones LR, Stark CE, Yassa MA, Bassett SS, Shelton AL, Gallagher M. 2012. Reduction of hippocampal hyperactivity improves cognition in amnestic mild cognitive impairment. Neuron 74:467–474. doi:10.1016/j.neuron.2012.03.023

Bankhead P, Loughrey MB, Fernández JA, Dombrowski Y, McArt DG, Dunne PD, McQuaid S, Gray RT, Murray LJ, Coleman HG, James JA, Salto-Tellez M, Hamilton PW. 2017. QuPath: Open source software for digital pathology image analysis. Sci Rep 7:16878. doi:10.1038/s41598-017-17204-5

Barker GRI, Banks PJ, Scott H, Ralph GS, Mitrophanous KA, Wong L-F, Bashir ZI, Uney JB, Warburton EC. 2017. Separate elements of episodic memory subserved by distinct hippocampal-prefrontal connections. Nat Neurosci 20:242–250. doi:10.1038/nn.4472

Berland C, Montalban E, Perrin E, Di Miceli M, Nakamura Y, Martinat M, Sullivan M, Davis XS, Shenasa MA, Martin C, Tolu S, Marti F, Caille S, Castel J, Perez S, Salinas CG, Morel C, Hecksher-Sørensen J, Cador M, Fioramonti X, Tschöp MH, Layé S, Venance L, Faure P, Hnasko TS, Small DM, Gangarossa G, Luquet S. 2020. Circulating triglycerides gate dopamine-associated behaviours through dopamine receptor type 2 (DRD2)-expressing neurons. Cell Metab 31:773–790.e11. doi:10.1016/j.cmet.2020.02.010

Biyong EF, Alfos S, Dumetz F, Helbling J-C, Aubert A, Brossaud J, Foury A, Moisan M-P, Layé S, Richard E, Patterson E, Murphy K, Rea K, Stanton C, Schellekens H, Cryan JF, Capuron L, Pallet V, Ferreira G. 2021. Dietary vitamin A supplementation prevents early obesogenic diet-induced microbiota, neuronal and cognitive alterations. Int J Obes 2005 45:588–598. doi:10.1038/s41366-020-00723-z

Boitard C, Cavaroc A, Sauvant J, Aubert A, Castanon N, Layé S, Ferreira G. 2014. Impairment of hippocampal-dependent memory induced by juvenile high-fat diet intake is associated with enhanced hippocampal inflammation in rats. Brain Behav Immun 40:9–17. doi:10.1016/j.bbi.2014.03.005

Boitard C, Etchamendy N, Sauvant J, Aubert A, Tronel S, Marighetto A, Layé S, Ferreira G. 2012. Juvenile, but not adult exposure to high-fat diet impairs relational memory and hippocampal neurogenesis in mice. Hippocampus 22:2095–2100. doi:10.1002/hipo.22032

Boitard C, Maroun M, Tantot F, Cavaroc A, Sauvant J, Marchand A, Layé S, Capuron L, Darnaudery M, Castanon N, Coutureau E, Vouimba R-M, Ferreira G. 2015. Juvenile obesity enhances emotional memory and amygdala plasticity through glucocorticoids. J Neurosci Off J Soc Neurosci 35:4092–4103. doi:10.1523/JNEUROSCI.3122-14.2015

Britt JP, Benaliouad F, McDevitt RA, Stuber GD, Wise RA, Bonci A. 2012. Synaptic and behavioral profile of multiple glutamatergic inputs to the nucleus accumbens. Neuron 76:790–803. doi:10.1016/j.neuron.2012.09.040

Carbone S, Ponzo OJ, Gobetto N, Samaniego YA, Reynoso R, Scacchi P, Moguilevsky JA, Cutrera R. 2013. Antiandrogenic effect of perinatal exposure to the endocrine disruptor di-(2-ethylhexyl) phthalate increases anxiety-like behavior in male rats during sexual maturation. Horm Behav 63:692–699. doi:10.1016/j.yhbeh.2013.01.006

Cenquizca LA, Swanson LW. 2007. Spatial organization of direct hippocampal field CA1 axonal projections to the rest of the cerebral cortex. Brain Res Rev 56:1–26. doi:10.1016/j.brainresrev.2007.05.002

Ciocchi S, Passecker J, Malagon-Vina H, Mikus N, Klausberger T. 2015. Brain computation. Selective information routing by ventral hippocampal CA1 projection neurons. Science 348:560–563. doi:10.1126/science.aaa3245

Décarie-Spain L, Sharma S, Hryhorczuk C, Issa-Garcia V, Barker PA, Arbour N, Alquier T, Fulton S. 2018. Nucleus accumbens inflammation mediates anxiodepressive behavior and compulsive sucrose seeking elicited by saturated dietary fat. Mol Metab 10:1–13. doi:10.1016/j.molmet.2018.01.018

Del Olmo N, Ruiz-Gayo M. 2018. Influence of High-Fat Diets Consumed During the Juvenile Period on Hippocampal Morphology and Function. Front Cell Neurosci 12:439. doi:10.3389/fncel.2018.00439

Ducrocq F, Hyde A, Fanet H, Oummadi A, Walle R, De Smedt-Peyrusse V, Layé S, Ferreira G, Trifilieff P, Vancassel S. 2019. Decrease in Operant Responding Under Obesogenic Diet Exposure is not Related to Deficits in Incentive or Hedonic Processes. Obesity 27:255–263. doi:10.1002/oby.22358

Ducrocq F, Walle R, Contini A, Oummadi A, Caraballo B, Veldt S van der, Boyer M-L, Aby F, Tolentino-Cortez T, Helbling J-C, Martine L, Grégoire S, Cabaret S, Vancassel S, Layé S, Kang JX, Fioramonti X, Berdeaux O, Barreda-Gómez G, Masson E, Ferreira G, Ma DWL, Bosch-Bouju C, Smedt-Peyrusse VD, Trifilieff P. 2020. Causal Link between n-3 Polyunsaturated Fatty Acid Deficiency and Motivation Deficits. Cell Metab 31:755–772.e7. doi:10.1016/j.cmet.2020.02.012

Francis H, Stevenson R. 2013. The longer-term impacts of Western diet on human cognition and the brain. Appetite 63:119–128. doi:10.1016/j.appet.2012.12.018

Friedman NP, Robbins TW. 2022. The role of prefrontal cortex in cognitive control and executive function. Neuropsychopharmacology 47:72–89. doi:10.1038/s41386-021-01132-0

Fulton S, Décarie-Spain L, Fioramonti X, Guiard B, Nakajima S. 2022. The menace of obesity to depression and anxiety prevalence. Trends Endocrinol Metab TEM 33:18–35. doi:10.1016/j.tem.2021.10.005

Gergues MM, Han KJ, Choi HS, Brown B, Clausing KJ, Turner VS, Vainchtein ID, Molofsky AV, Kheirbek MA. 2020. Circuit and molecular architecture of a ventral hippocampal network. Nat Neurosci 23:1444–1452. doi:10.1038/s41593-020-0705-8

Glushchak K, Ficarro A, Schoenfeld TJ. 2021. High-fat diet and acute stress have different effects on object preference tests in rats during adolescence and adulthood. Behav Brain Res 399:112993. doi:10.1016/j.bbr.2020.112993

Hwang L-L, Wang C-H, Li T-L, Chang S-D, Lin L-C, Chen C-P, Chen C-T, Liang K-C, Ho I-K, Yang W-S, Chiou L-C. 2010. Sex differences in high-fat diet-induced obesity, metabolic alterations and learning, and synaptic plasticity deficits in mice. Obes Silver Spring Md 18:463–469. doi:10.1038/oby.2009.273

Kerfoot EC, Williams CL. 2018. Contributions of the Nucleus Accumbens Shell in Mediating the Enhancement in Memory Following Noradrenergic Activation of Either the Amygdala or Hippocampus. Front Pharmacol 9:47. doi:10.3389/fphar.2018.00047

Khan NA, Baym CL, Monti JM, Raine LB, Drollette ES, Scudder MR, Moore RD, Kramer AF, Hillman CH, Cohen NJ. 2015. Central Adiposity is Negatively Associated with Hippocampal-Dependent Relational Memory among Overweight and Obese Children. J Pediatr 166:302–308.e1. doi:10.1016/j.jpeds.2014.10.008

Khazen T, Hatoum OA, Ferreira G, Maroun M. 2019. Acute exposure to a high-fat diet in juvenile male rats disrupts hippocampal-dependent memory and plasticity through glucocorticoids. Sci Rep 9:12270. doi:10.1038/s41598-019-48800-2

Labouesse MA, Lassalle O, Richetto J, Iafrati J, Weber-Stadlbauer U, Notter T, Gschwind T, Pujadas L, Soriano E, Reichelt AC, Labouesse C, Langhans W, Chavis P, Meyer U. 2017. Hypervulnerability of the adolescent prefrontal cortex to nutritional stress via reelin deficiency. Mol Psychiatry 22:961–971. doi:10.1038/mp.2016.193

Labouesse MA, Stadlbauer U, Langhans W, Meyer U. 2013. Chronic high fat diet consumption impairs sensorimotor gating in mice. Psychoneuroendocrinology 38:2562–2574. doi:10.1016/j.psyneuen.2013.06.003

Leger M, Quiedeville A, Bouet V, Haelewyn B, Boulouard M, Schumann-Bard P, Freret T. 2013. Object recognition test in mice. Nat Protoc 8:2531–2537. doi:10.1038/nprot.2013.155

Li M, Long C, Yang L. 2015. Hippocampal-prefrontal circuit and disrupted functional connectivity in psychiatric and neurodegenerative disorders. BioMed Res Int 2015:810548. doi:10.1155/2015/810548

Liu X, Carter AG. 2018. Ventral Hippocampal Inputs Preferentially Drive Corticocortical Neurons in the Infralimbic Prefrontal Cortex. J Neurosci 38:7351–7363. doi:10.1523/JNEUROSCI.0378-18.2018

Martin AA, Davidson TL. 2014. Human Cognitive Function and the Obesogenic Environment. Physiol Behav 0:185–193. doi:10.1016/j.physbeh.2014.02.062

Moreno-Castilla P, Guzman-Ramos K, Bermudez-Rattoni F. 2018. Object Recognition and Object Location Recognition Memory – The Role of Dopamine and NoradrenalineHandbook of Behavioral Neuroscience. Elsevier. pp. 403–413. doi:10.1016/B978-0-12-812012-5.00028-8

Morici JF, Weisstaub NV, Zold CL. 2022. Hippocampal-medial prefrontal cortex network dynamics predict performance during retrieval in a context-guided object memory task. Proc Natl Acad Sci 119:e2203024119. doi:10.1073/pnas.2203024119

Morin J-P, Rodríguez-Durán LF, Guzmán-Ramos K, Perez-Cruz C, Ferreira G, Diaz-Cintra S, Pacheco-López G. 2017. Palatable Hyper-Caloric Foods Impact on Neuronal Plasticity. Front Behav Neurosci 11:19. doi:10.3389/fnbeh.2017.00019

Murray S, Chen EY. 2019. Examining Adolescence as a Sensitive Period for High-Fat, High-Sugar Diet Exposure: A Systematic Review of the Animal Literature. Front Neurosci 13:1108. doi:10.3389/fnins.2019.01108

Naneix F, Bakoyiannis I, Santoyo-Zedillo M, Bosch-Bouju C, Pacheco-Lopez G, Coutureau E, Ferreira G, OBETEEN Consortium. 2021. Chemogenetic silencing of hippocampus and amygdala reveals a double dissociation in periadolescent obesogenic diet-induced memory alterations. Neurobiol Learn Mem 178:107354. doi:10.1016/j.nlm.2020.107354

Naneix F, Tantot F, Glangetas C, Kaufling J, Janthakhin Y, Boitard C, De Smedt-Peyrusse V, Pape JR, Vancassel S, Trifilieff P, Georges F, Coutureau E, Ferreira G. 2017. Impact of Early Consumption of High-Fat Diet on the Mesolimbic Dopaminergic System. eNeuro 4:ENEURO.0120-17.2017. doi:10.1523/ENEURO.0120-17.2017

Nelson AJD, Cooper MT, Thur KE, Marsden CA, Cassaday HJ. 2011. The Effect of Catecholaminergic Depletion Within the Prelimbic and Infralimbic Medial Prefrontal Cortex on Recognition Memory for Recency, Location, and Objects. Behav Neurosci 125:396–403. doi:10.1037/a0023337

Nyaradi A, Foster JK, Hickling S, Li J, Ambrosini GL, Jacques A, Oddy WH. 2014. Prospective associations between dietary patterns and cognitive performance during adolescence. J Child Psychol Psychiatry 55:1017–1024. doi:10.1111/jcpp.12209

Ogden CL, Carroll MD, Lawman HG, Fryar CD, Kruszon-Moran D, Kit BK, Flegal KM. 2016. Trends in Obesity Prevalence Among Children and Adolescents in the United States, 1988-1994 Through 2013-2014. JAMA 315:2292–2299. doi:10.1001/jama.2016.6361

Øverby NC, Lüdemann E, Høigaard R. 2013. Self-reported learning difficulties and dietary intake in Norwegian adolescents. Scand J Public Health 41:754–760. doi:10.1177/1403494813487449

Padilla-Coreano N, Bolkan SS, Pierce GM, Blackman DR, Hardin WD, Garcia-Garcia AL, Spellman TJ, Gordon JA. 2016. Direct Ventral Hippocampal-Prefrontal Input Is Required for Anxiety-Related Neural Activity and Behavior. Neuron 89:857–866. doi:10.1016/j.neuron.2016.01.011

Palmer BF, Clegg DJ. 2015. The sexual dimorphism of obesity. Mol Cell Endocrinol 0:113–119. doi:10.1016/j.mce.2014.11.029

Parfitt GM, Nguyen R, Bang JY, Aqrabawi AJ, Tran MM, Seo DK, Richards BA, Kim JC. 2017. Bidirectional Control of Anxiety-Related Behaviors in Mice: Role of Inputs Arising from the Ventral Hippocampus to the Lateral Septum and Medial Prefrontal Cortex. Neuropsychopharmacol Off Publ Am Coll Neuropsychopharmacol 42:1715–1728. doi:10.1038/npp.2017.56

Paxinos G, Franklin K. 2004. Paxinos and Franklin’s the Mouse Brain in Stereotaxic Coordinates - 5th Edition. https://www.elsevier.com/books/paxinos-and-franklins-the-mouse-brain-in-stereotaxic-coordinates/paxinos/978-0-12-816157-9

Phillips ML, Robinson HA, Pozzo-Miller L. 2019. Ventral hippocampal projections to the medial prefrontal cortex regulate social memory. eLife 8:e44182. doi:10.7554/eLife.44182

Reichelt AC, Gibson GD, Abbott KN, Hare DJ. 2019. A high-fat high-sugar diet in adolescent rats impairs social memory and alters chemical markers characteristic of atypical neuroplasticity and parvalbumin interneuron depletion in the medial prefrontal cortex. Food Funct 10:1985–1998. doi:10.1039/c8fo02118j

Roozendaal B, Quervain DJ-F de, Ferry B, Setlow B, McGaugh JL. 2001. Basolateral Amygdala–Nucleus Accumbens Interactions in Mediating Glucocorticoid Enhancement of Memory Consolidation. J Neurosci 21:2518–2525. doi:10.1523/JNEUROSCI.21-07-02518.2001

Sargolini F, Roullet P, Oliverio A, Mele A. 2003. Effects of intra-accumbens focal administrations of glutamate antagonists on object recognition memory in mice. Behav Brain Res 138:153–163. doi:10.1016/s0166-4328(02)00238-3

Sellbom KS, Gunstad J. 2012. Cognitive function and decline in obesity. J Alzheimers Dis JAD 30 Suppl 2:S89–95. doi:10.3233/JAD-2011-111073

Spear LP. 2000. The adolescent brain and age-related behavioral manifestations. Neurosci Biobehav Rev 24:417–463. doi:10.1016/s0149-7634(00)00014-2

Sui SX, Pasco JA. 2020. Obesity and Brain Function: The Brain-Body Crosstalk. Med Kaunas Lith 56:E499. doi:10.3390/medicina56100499

Tsan L, Décarie-Spain L, Noble EE, Kanoski SE. 2021. Western Diet Consumption During Development: Setting the Stage for Neurocognitive Dysfunction. Front Neurosci 15:632312. doi:10.3389/fnins.2021.632312

Valladolid-Acebes I, Fole A, Martín M, Morales L, Cano MV, Ruiz-Gayo M, Del Olmo N. 2013. Spatial memory impairment and changes in hippocampal morphology are triggered by high-fat diets in adolescent mice. Is there a role of leptin? Neurobiol Learn Mem 106:18–25. doi:10.1016/j.nlm.2013.06.012

Vega-Torres JD, Haddad E, Lee JB, Kalyan-Masih P, Maldonado George WI, López Pérez L, Piñero Vázquez DM, Arroyo Torres Y, Santiago Santana JM, Obenaus A, Figueroa JD. 2018. Exposure to an obesogenic diet during adolescence leads to abnormal maturation of neural and behavioral substrates underpinning fear and anxiety. Brain Behav Immun 70:96–117. doi:10.1016/j.bbi.2018.01.011

Vouimba R-M, Bakoyiannis I, Ducourneau E-G, Maroun M, Ferreira G. 2021. Bidirectional modulation of hippocampal and amygdala synaptic plasticity by post-weaning obesogenic diet intake in male rats: Influence of the duration of diet exposure. Hippocampus 31:117–121. doi:10.1002/hipo.23278

Wang C, Niu R, Zhu Y, Han H, Luo G, Zhou B, Wang J. 2014. Changes in memory and synaptic plasticity induced in male rats after maternal exposure to bisphenol A. Toxicology 322:51–60. doi:10.1016/j.tox.2014.05.001

Yaseen A, Shrivastava K, Zuri Z, Hatoum OA, Maroun M. 2019. Prefrontal Oxytocin is Involved in Impairments in Prefrontal Plasticity and Social Memory Following Acute Exposure to High Fat Diet in Juvenile Animals. Cereb Cortex N Y N 1991 29:1900–1909. doi:10.1093/cercor/bhy070

Yassa MA, Lacy JW, Stark SM, Albert MS, Gallagher M, Stark CEL. 2011. Pattern separation deficits associated with increased hippocampal CA3 and dentate gyrus activity in nondemented older adults. Hippocampus 21:968–979. doi:10.1002/hipo.20808

Yeomans MR. 2017. Adverse effects of consuming high fat-sugar diets on cognition: implications for understanding obesity. Proc Nutr Soc 76:455–465. doi:10.1017/S0029665117000805

